# Uniform Genomic Data Analysis in the NCI Genomic Data Commons

**DOI:** 10.1101/788919

**Authors:** Zhenyu Zhang, Kyle Hernandez, Jeremiah Savage, Shenglai Li, Dan Miller, Stuti Agrawal, Francisco Ortuno, Lou Staudt, Allison Heath, Robert L. Grossman

## Abstract

The goal of the National Cancer Institute (NCI) Genomic Data Commons (GDC) is to provide the cancer research community with a data repository of uniformly processed genomic and associated clinical data that enables data sharing and collaborative analysis in the support of precision medicine. The initial GDC dataset include genomic, epigenomic, proteomic, clinical and other data from the NCI TCGA and TARGET programs. Data production for the GDC started in June, 2015 using an OpenStack-based private cloud. By June of 2016, the GDC had analyzed more than 50,000 raw sequencing data inputs, as well as multiple other data types. Using the latest human genome reference build GRCh38, the GDC generated a variety of data types from aligned reads to somatic mutations, gene expression, miRNA expression, DNA methylation status, and copy number variation. In this paper, we describe the pipelines and workflows used to process and harmonize the data in the GDC. The generated data, as well as the original input files from TCGA and TARGET, are available for download and exploratory analysis at the GDC Data Portal and Legacy Archive (https://gdc.cancer.gov/).

## Introduction

The National Cancer Institute’s (NCI) Genomic Data Commons (GDC) ^1^ currently contains NCI-generated data from some of the largest and most comprehensive cancer genomic datasets, including The Cancer Genome Atlas (TCGA, https://cancergenome.nih.gov/) and Therapeutically Applicable Research to Generate Effective Therapies (TARGET, https://ocg.cancer.gov/programs/target). Each of these projects contains a variety of processed and unprocessed molecular data types, including genomics, epigenomics, proteomics, imaging, clinical, and others.

These data, as well as data from future projects, are often generated using different methods, so that joint analysis of multiple datasets are often confounded by batch effects. Based on the lessons learned from TCGA, one of the major goals of the GDC is to create a uniform set of molecular datasets that minimize batch effects due to differences in reference genomes, gene models, analytical algorithms, and processing pipelines. In the GDC, this process is called harmonization and is broken up into two stages: alignment and higher level data generation. In the alignment stage, reads from Next Generation Sequencing (NGS) data are extracted and aligned, or re-aligned, to a single human reference genome sequence using a single pipeline for that particular data type. In the second stage, different higher level data generation pipelines utilize the GDC aligned data to derive summary results, such as somatic mutations or gene expression.

Data production started with processing of Whole Exome Sequencing (WXS) alignment and somatic mutation calling, Whole Genome Sequencing (WGS) alignment, mRNA-Seq alignment and gene/exon level quantification, miRNA-Seq alignment and quantification, TCGA Affymetrix Genome-Wide Human SNP Array 6.0 array copy number segmentation, and Illumina Infinium HumanMethylation450/27 methylation array re-annotation. By the end of May 2016, we had successfully processed more than 56,168 BAM files and FASTQ bundles, including 27,594 WXS, 4691 WGS, 11,969 RNA-Seq and 11,914 miRNA-Seq, from 12,487 patients with a total input file size of 1001.7 TB. From GDC harmonized BAMs, more than 100,000 higher level derived data files have been generated. A total of more than 1400 TB of generated data, including intermediate files, have been produced by GDC data processing pipelines, and stored in GDC storage. Many of them have been imported into the GDC database (described in a companion submission), and have become queryable and downloadable from the GDC Data Portal (https://portal.gdc.cancer.gov/) and API.

## Results

### 1 Reference Genome and Gene Model

The GDC chose GRCh38 as the reference human genome build for all data analyses, because of its improved coverage and accuracy over the previous major build GRCh37 ^2^. The GRCh38 major human genome assembly was released by GRC (Genome Reference Consortium, https://www.ncbi.nlm.nih.gov/grc/) on Dec 2013 with GenBank assembly accession GCA_000001405.15. The complete assembly downloaded from NCBI contains 456 sequences, including 25 continuous chromosomal and mitochondrial sequences, 42 unlocalized scaffolds, 127 unplaced scaffolds, 261 alternative scaffolds, and 1 EBV decoy sequence.

At the time of GDC pipeline development, there was a lack of tool support for alternative scaffolds analysis. As a result, the GDC excluded alternative contigs from the GDC reference sequence, and used GCA_000001405.15_GRCh38_no_alt_analysis_set from the NCBI ftp site (ftp://ftp.ncbi.nlm.nih.gov/genomes/all/GCA/000/001/405/GCA_000001405.15_GRCh38/seqs_for_alignment_pipelines.ucsc_ids/GCA_000001405.15_GRCh38_no_alt_analysis_set.fna.gz, see description README_analysis_sets.txt in the same directory).

In addition to the continuous chromosomal and mitochondrial sequences, unlocalized and unplaced scaffolds described above, the GDC reference sequence also includes 2385 human decoy sequences, together named hs38d1 with GenBank assembly accession GCA_000786075.2. These human decoy sequences are additional unlocalized sequences not officially recognized by the Genome Reference Consortium at the time of reference release, and having them in the GDC reference helps to reduce false alignments of such reads in other regions ^3^.

Similar to the idea of improving alignment quality with human decoy sequence, we collected genomic sequences from 10 types (200 subtypes) of cancer related viruses in the reference genome, together called “virus decoy” as a collection. The additional benefit of having such virus sequences in the reference genome is to give users the opportunity to directly identify virus genome derived reads. These viruses include Human Cytomegalovirus (CMV, HHV-5), Epstein-Barr virus (EBV, HHV-4), Hepatitis B virus (HBV), Hepatitis C virus (HCV), Human immunodeficiency virus (HIV), Human herpesvirus 8 (KSHV, HHV-8), Human T-lymphotropic virus 1 (HTLV-1), Merkel cell polyomavirus (MCV), Simian vacuolating virus 40 (SV40) and Human papillomavirus (HPV) (Table 1). All HPV sequences were obtained from The PapillomaVirus Episteme (PaVE, http://pave.niaid.nih.gov/) instead of NCBI, because this group updated the sequences as new information is confirmed ^4^. The GDC reference including all human and viral decoys can be downloaded from the GDC at the following link (https://api.gdc.cancer.gov/data/62f23fad-0f24-43fb-8844-990d531947cf).

**Table 1.** Virus Sequences in GDC Reference Genome (Full table is provided in a separate Excel spreadsheet)

| Contig_Name | Virus_Name | GenBank_Accession | PaVE_ID | Revised_Seq |
| --- | --- | --- | --- | --- |
| CMV | Human Cytomegalovirus, Human herpes virus 5 | AY446894.2 |  |  |
| EBV | Epstein-Barr virus, , Human herpes virus 4 | AJ507799.2 |  |  |
| HBV | Hepatitis B | X04615.1 |  |  |
| HCV-1 | Hepatitis C | AF009606.1 |  |  |
| HCV-2 | Hepatitis C | AF177036.1 |  |  |
| HIV-1 | Human immunodeficiency virus 1 | AF033819.3 |  |  |
| HIV-2 | Human immunodeficiency virus 2 | M30502.1 |  |  |
| KSHV | Kaposi's sarcoma-associated herpesvirus, Human herpes virus 8 | AF148805.2 |  |  |
| HTLV-1 | Human T-lymphotropic virus 1 | AF033817.1 |  |  |
| MCV | Merkel cell polyomavirus | HM011556.1 |  |  |
| SV40 | Simian vacuolating virus 40 | J02400.1 |  |  |
| HPV16 | Human papillomavirus 16 | K02718 | HPV16REF | Revised |
| HPV18 | Human papillomavirus 18 | X05015 | HPV18REF | Revised |

\* All HPV sequences are retrieved from the Papillomavirus Knowledge Source (PaVE) <sup>4</sup> as of May 2015. Genomic sequences of 18 HPV subtypes were corrected by re-sequencing or efforts to maintain integrity of characterized open-reading frames <sup>4</sup>, and thus not are consistent with sequences in the GenBank.

For variant annotation and RNA-Seq alignment/quantification, we use Human GENCODE release version 22 as the default GRCh38 gene model ^4,5^. This version contains 60,483 genes, including 19,814 protein-coding genes, 15,900 long non-coding RNA genes, 9894 small non-coding RNA genes, and many other types, such as pseudogenes. For miRNA-Seq annotation and qualifications, GDC uses miRBase version 21 ^6^.

### 2 Reproducibility of analysis

All major GDC data production pipelines are written in the Common Workflow Language (CWL, https://www.commonwl.org/). In each workflow, a main CWL file describes how tools and sub-workflows, also written in CWL, can be used in clearly defined steps. All major tools have been containerized using Docker containers to support reproducibility and portability of the workflows. The GDC will be redistributing the main GDC workflows to the research community to support reproducible research.

### 3 DNA-Seq Alignment, Mutation Calling, Annotation, and Somatic Variant Aggregation

The DNA-Seq alignment process includes initial alignment and a post-alignment optimization process. In the initial alignment step, reads are mapped to GRCh38 using BWA, followed by BAM sorting, merging of read group BAMs into a single BAM, and then duplicate-marking using Picard. When the read length of a read group is larger than 70 bps, BWA MEM ^7^ is used for alignment; or otherwise, BWA Aln ^7,8^ is used.

After intial alignment of WXS data, we follow GATK Best Practice (https://software.broadinstitute.org/gatk/best-practices/) to process all BAMs from the same patient together for a post-alignment optimization process called “co-cleaning” in which includes GATK IndelRealigner and BaseQualityScoreRecalibration (BQSR). IndelRealigner performs local realignment to further improve mapping quality acrossing all reads at loci close to indels, and BQSR detects and fixes systematic errors made by the sequencer when it estimates the quality score of each base call ^9–11^. For post-alignment optimization of WGS data, we only employ BQSR, but not IndelRealigner.

GDC uses four somatic variant callers, MuSE ^12^, MuTect2 ^11^, VarScan2 ^13^, and SomaticSniper^14^. MuSE and SomaticSniper only detect point mutations, while MuTect2 and VarScan2 can detect both point mutations and small insertions and deletions (INDELs). The GDC provides these point mutations and small INDELs together as Simple Nucleotide Variants (SNV). INDEL calls from VarScan2 were not included in the initial GDC data release, but is included since release 10.

Artificial chimera reads can form during the Multiple Displacement Amplification reaction ^15^, such as the one used to generate the WGA libraries by REPLI-g. This phenomenon is most obvious in MuTect2 calls shown as abnormally high INDEL/SNP ratio, and we observed that most of such INDELs are primarily supported by soft-clipped reads. In order to increase specificity, we re-analyzed all TCGA tumor WGA samples with MuTect2 option –dontUseSoftClippedBases, and successfully removed most of these false-positive INDELs. However, as these artifacts were introduced during library preparation, they could also exist in MuTect2 SNPs, as well as variants called by other tools, in a smaller scale. We recommend users to consume these WGA calls with care.

Raw Somatic Variant Call Format files (VCFs) generated from each pipeline are further processed by caller-specific filters to tag low quality variants in the FILTER column in VCF. For SomaticSniper, variants with Somatic Score (SSC) < 25 are removed. These VCFs are then annotated using Variant Effect Predictor (VEP) ^16^ to generate Annotated Somatic VCFs.

To allow easier investigation of variants and annotations, the VCFs are transformed into project-level tab-delimited Mutation Annotation Format (MAF) files ^17^. This is done with a custom tool based upon VCF2MAF (https://github.com/mskcc/vcf2maf) from Memorial Sloan Kettering Cancer Center. VCF and MAF files may contain germline variants and therefore all VCFs and MAFs described above are only available as controlled-access files. To access controlled-access files in the GDC, users must obtain approval from the Data Access Committee through dbGaP (database of Genotypes and Phenotypes) ^18^.

We have also created open-access MAF files by applying stringent criteria to remove potential germline variants. These open-access MAF files are used to support variant visualization and simpler data sharing. We didn’t produce open-access MAFs for the TARGET program because of privacy concerns of child sample donors. Mutation loads of point mutations and INDELs from both open-access (public) and controlled-access (protected) MAFs of all TCGA projects are displayed in Figure 1. Of note, this germline-masking process is so stringent that some real somatic variants, for example somatic variants in areas of low sequencing coverage in the paired normal samples, may have been removed from open-access MAF. We encourage users to explore controlled-access MAFs to view additional variants.

**Figure 1.**
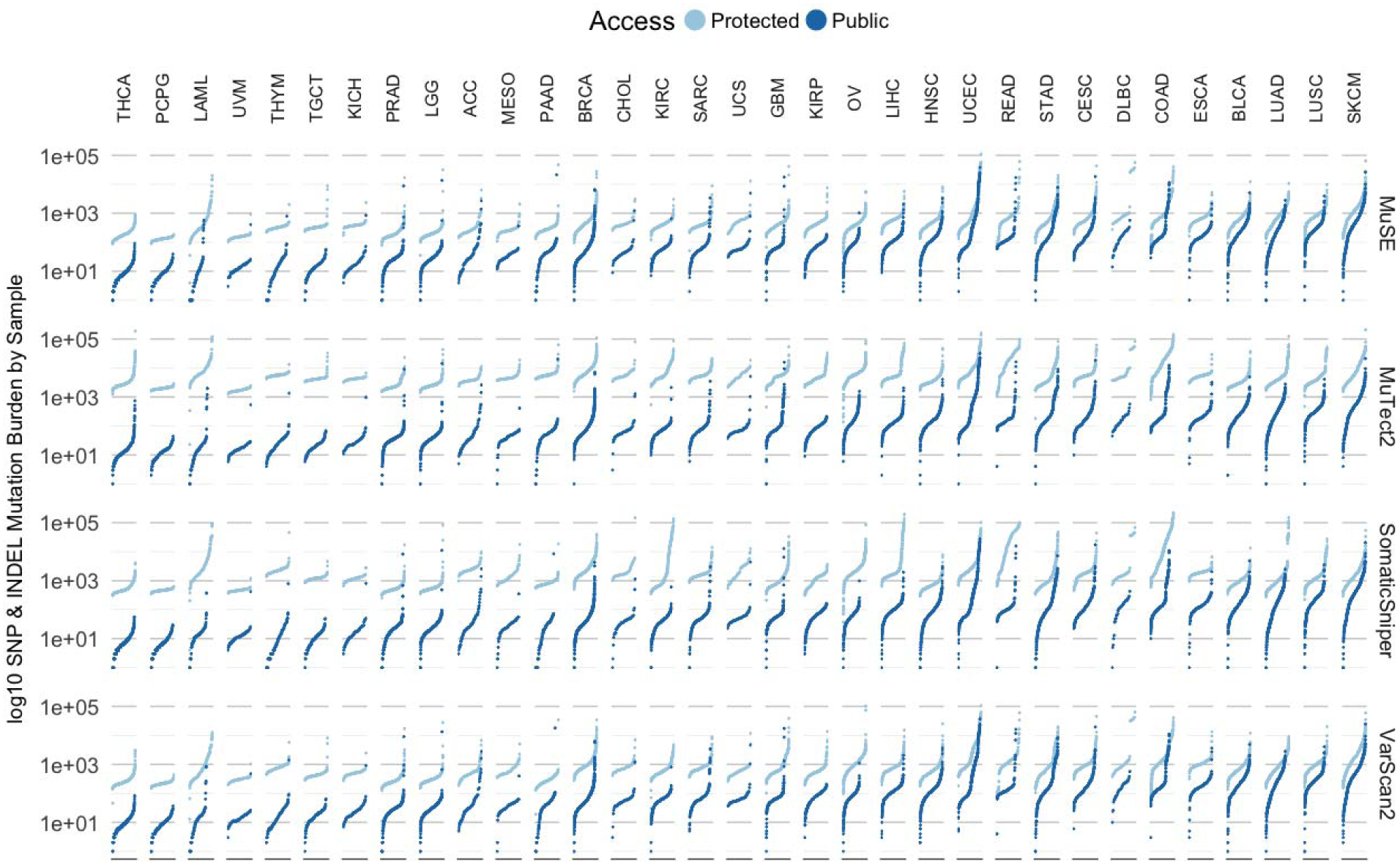
Mutation Loads of TCGA Projects. GDC-detected somatic variants per sample are displayed by each pipeline (rows), and grouped in each project (columns). Combined counts of point mutations (SNP) and INDELs of either public MAF or protected MAF are plotted in separate colors.

### 4 Quality Assessment of GDC Somatic Variants

Somatic variant detection is still in a stage of rapid algorithm development, and no single caller is superior to others in every respect ^19–22^. As shown in Figure 2A, a direct comparison of SNV calls from different callers show significant overlaps, but also many tool-specific calls. Here, we only compared high quality variants, which means they do not have a non-PASS caller assigned FILTER value or GDC assigned GDC_FILTER values in the somatic MAF files of data release version 10. In summary, 56.0% of the clean variants have been identified by all four callers, 15.1% by three callers, 14.0% by two callers, and 14.9% called by only one caller. Among all four somatic callers, MuTect2 has detected the highest number of unique calls, while SomaticSniper has the least. Additional efforts are needed to examine the validity of such calls, especially those not called by all pipelines, in order to evaluate the performance of each tool, and hopefully lead to machine-learning algorithms that help to unify caller outputs. Please note that these results are only relevant to the GDC implementation of these pipelines and filtering strategy.

**Figure 2.**
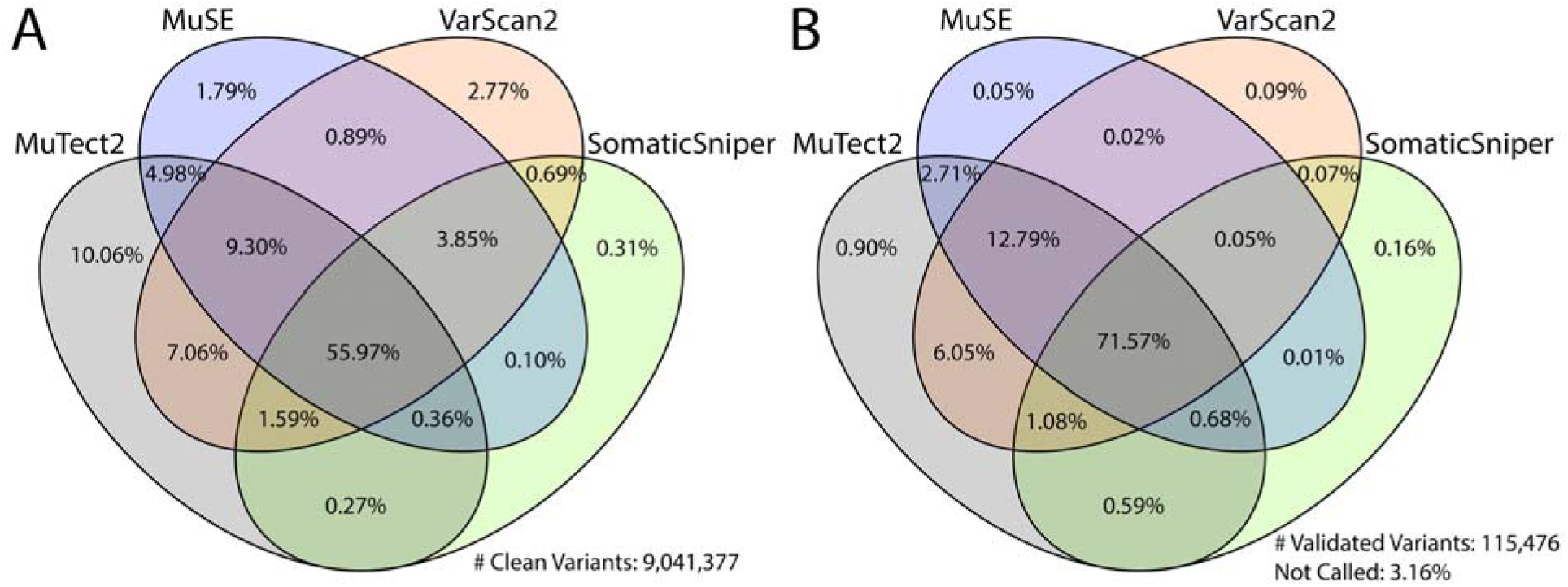
Comparison of GDC Somatic Variant Caller Pipelines. The Venn Diagram on the left (A) shows the overlap among four GDC somatic callers. Among all clean variants, 56.0% have been identified by all four callers, 15.1% by three callers, 14.0% by two callers, and 14.9% by only one caller. The Venn Diagram on the right (B) shows recall rate of validated TCGA variants by GDC somatic callers. Among 115,476 TCGA validated variants collected, 3.2% are not recalled by any of the GDC pipelines; 1.2% are recalled by only one pipeline; 9.4% are recalled by two pipelines; 14.6% are recalled by three pipelines; and 71.6% are recalled by all four GDC pipelines.

Evaluation of somatic variant callers often requires comparison with a so-called gold standard dataset that has been extensively sequenced using multiple independent methods ^22^, or using a simulated dataset. In a previous effort to evaluate quality of somatic variant callers, TCGA re-sequenced regions of many called variants using orthogonal sequence technologies, such as Sanger Sequencing, and therefore have produced a set of validated variants that GDC can use to evaluate False-Negatives (FN) in current GDC callers. However, no validation experiments were designed for GDC-called variants, thus False-Positives (FP) will not be evaluated in this paper.

We extracted and lifted over all TCGA validation information from 196 MAFs available on May 2016 to GRCh38 coordinates, and selected mutation calls from the same tumor samples that GDC has also successfully analyzed with all four pipelines. This results in 1,911 unique tumor samples and 115,476 validated variants, across 13 TCGA projects (BLCA, BRCA, CESC, COAD, GBM, KIRC, LAML, OV, PAAD, READ, SARC, THYM and UCS). Comparison to GDC called somatic variants of the sample tumor sample shows that only 3.2% of TCGA validated variants are not re-discovered by any of the GDC pipelines (Figure 2B). This number further decreases to 1.6% if we only compare variants from exactly the same tumor and normal aliquot pairs. 95.6% of the validated variants are called by at least two GDC pipelines; 86.2% by at least three pipelines; and 71.6% called by all four GDC pipelines.

To further investigate the impact of cancer type or project to each caller, we analyzed the validated variant recall rate for each pipeline (Figure 3). In most of the cancer types, MuTect2, MuSE and VarScan2 show good performance, while SomaticSniper typically recalls less. In particular, SomaticSniper displays a very low average recall rate of less than 50% in PAAD (Pancreatic Adenocarcinoma). Interestingly, SomaticSniper has the best recall rate for LAML. The algorithm of SomaticSniper may have been designed to better tolerate high-level tumor contaminations in germline control samples, which often exists as infiltration of liquid tumor cells in skin or buccal swabs of LAML patients.

**Figure 3.**
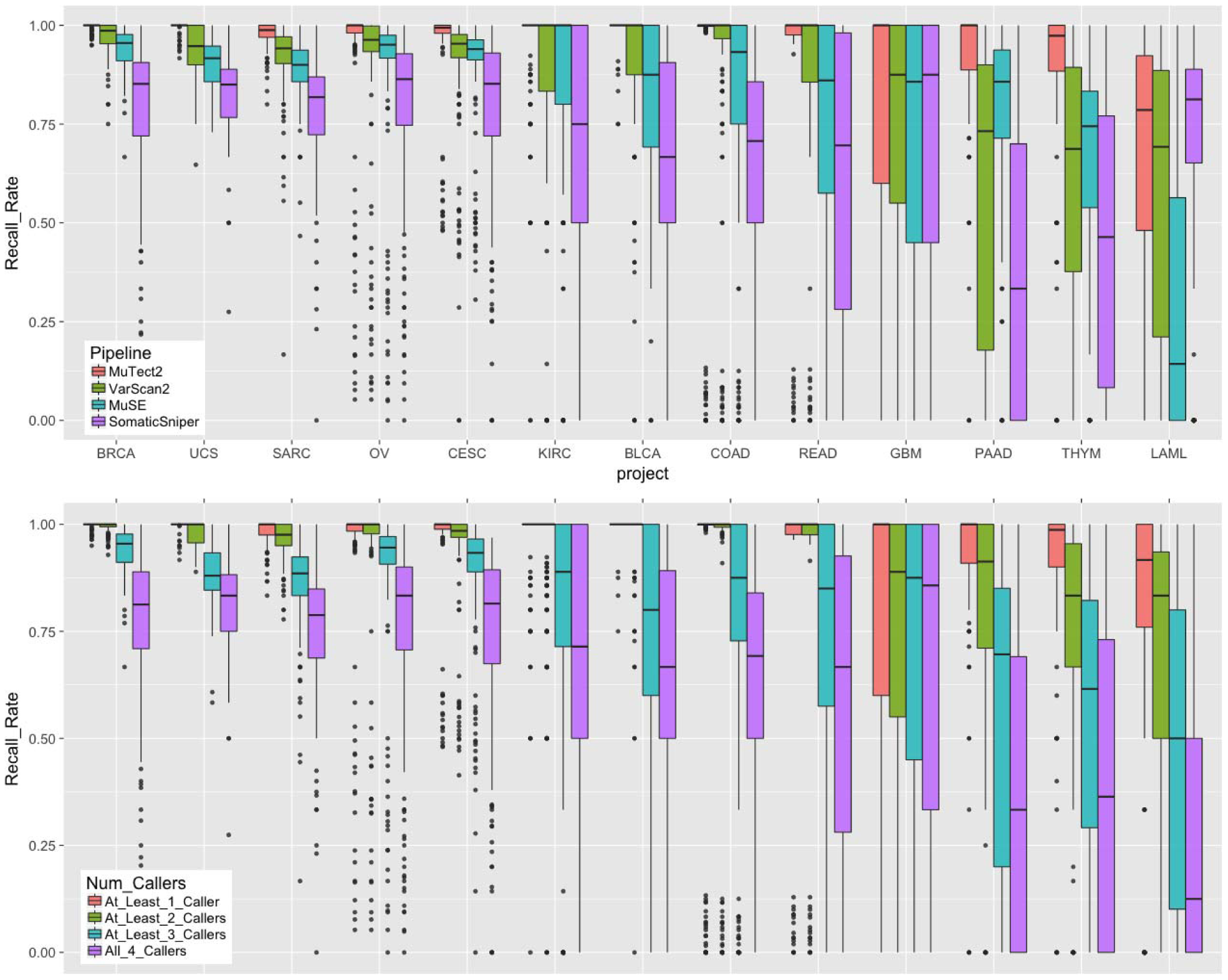
Recall Rate of TCGA Validated Variants by Project. Top: Boxplots of recall rate of TCGA validated variants by 13 projects and four GDC somatic variant calling pipelines. Each dot represents a unique tumor sample. Projects are ordered by decreasing average recall rate from left to right. Bottom: Boxplots of recall rate of TCGA validated variants by number of pipelines combined.

### 5 RNA-Seq Data Processing and Quality Assessment

RNA-Seq analysis by the GDC makes use of a modified workflow (https://github.com/ucscCancer/icgc_rnaseq_align) created by the International Cancer Genome Consortium (ICGC). Input RNA-Seq FASTQ files were aligned to GRCh38 using the STAR 2-pass method ^23^, and quantified using HTSeq ^24^ and DEXSeq ^25^. The GDC uses GENCODE v22 as the default gene model for both mRNA-Seq alignment and quantification. DEXSeq exon-level quantification results have not yet been imported into GDC Data Portal at the time of publication.

Gene expression is measured using HTSeq. In this pipeline, only reads or read pairs that can be uniquely assigned to a gene are counted. HTSeq supports mapping of both stranded (either forward or reverse) and unstranded libraries; however, the read assignment is different between these two different modes resulting in an inability to compare across these library types. Because the GDC emphasizes data comparability across projects, we have run HTSeq on all projects as if they were unstranded libraries.

For convenience to the scientific community, the GDC also produces gene level quantification in the units of FPKM (Fragments Per Kilobase of transcript per Million mapped reads) ^26^ and FPKM-UQ (Upper-Quartile Normalized FPKM) ^26,27^. The definition of these units are described in the Materials and Methods section. Note that the denominators of such normalizations are read counts of all the protein coding genes, instead of all genes. If users are interested in a different set of genes, they are encouraged to perform a normalization based on the genes they are working on, or to use a more sophisticated method, such as DESeq ^28^ or EdgeR ^29^.

We also compared GDC FPKM-UQ expression data to the original TCGA upper-quartile normalized RSEM expression values using Spearman correlation. This comparison was performed over 10,243 shared aliquots and 18,038 shared genes in both dataset. The average correlation between the same sample from two datasets (Figure 4. Top) is 94.4%, and the majority of the samples have correlation higher than 90%.

**Figure 4.**
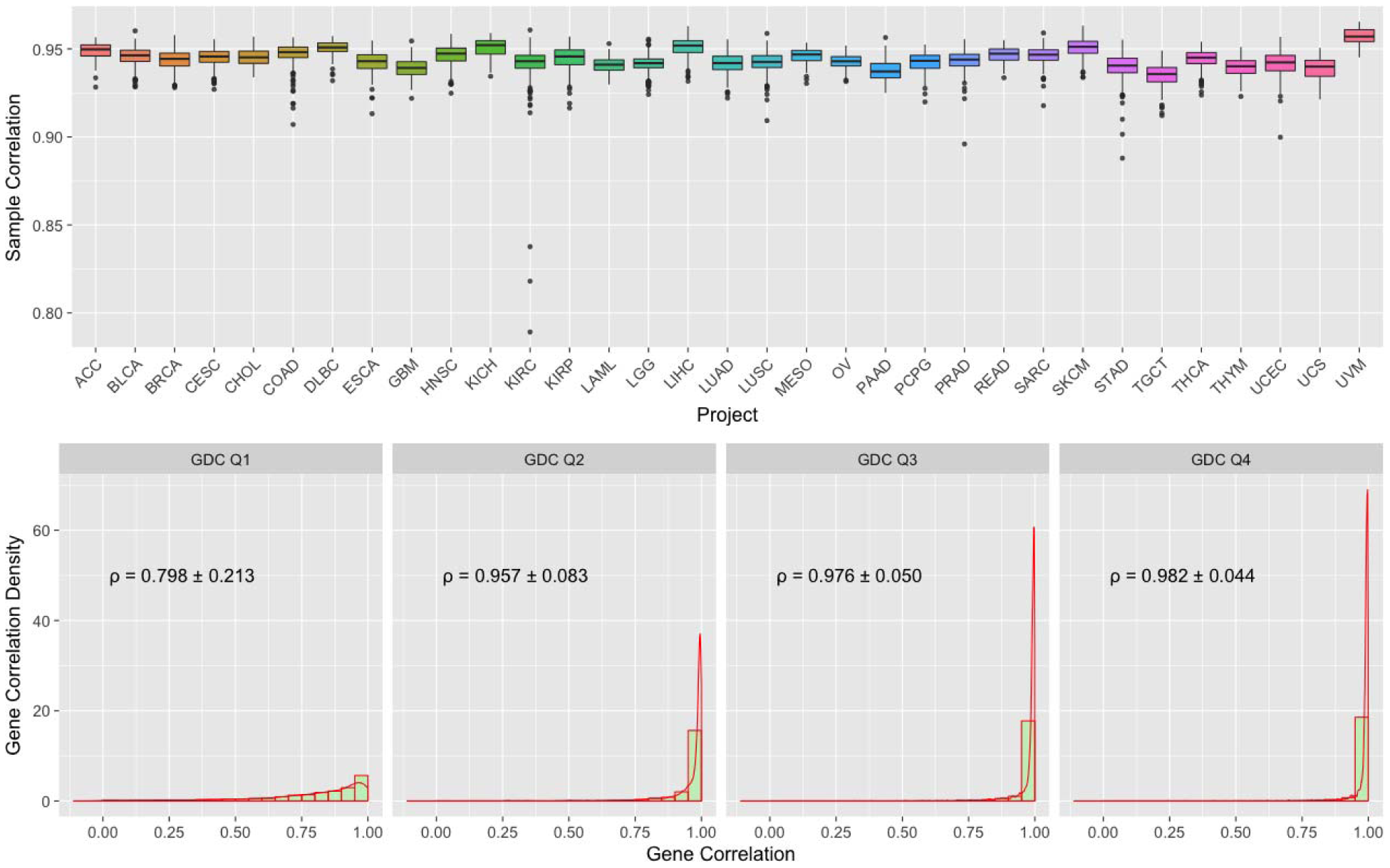
Boxplots of Spearman Correlation between GDC and TCGA mRNA Expression. Top: Boxplots of Sample to Sample Correlation between GDC and TCGA by Project. Bottom: Combined Boxplots and Density Plots of Gene to Gene Correlation between GDC and TCGA. All genes are categorized by four GDC groups (Q1-4) based on their average expression values. Mean and standard deviation of gene to gene Spearman’s correlations are calculated by these four groups.

We can also measure the relative expression of the same gene among different samples. The average correlation between the same gene from two datasets is 92.9%. We suspected that most of the deviation is from sporadic low-level expressed genes. To address this concern, we further categorized genes into 4 quartile groups (Q1 to Q4) based on their average expression values in the GDC, and then examined gene level correlations within each of these 4 groups with TCGA results (Figure 4. Bottom). We observed an average correlation of 98% in high-level expressed Q3 and Q4 groups and much lower values in Q1 and Q2.

### 6. miRNA Data Processing and Quality Assessment

The GDC miRNA quantification analysis workflow is based on the profiling pipeline that was developed by the British Columbia Genome Sciences Centre ^30^. After realignment of miRNA-Seq reads to GRCh38 using BWA Aln, the profiling pipeline generates TCGA-formatted miRNA gene expression and isoform expression results by comparing the individual reads to sequence feature annotations in miRBase v.21 ^6^. Of note, however, the tool only annotates those reads that have an exact match with known miRNAs in miRBase and therefore does not identify novel miRNA or transcript with mutations.

We compared TPM (Transcripts Per Kilobase Million) normalized miRNA gene expression from GDC GRCh38 pipeline to the original TCGA Hg19 pipeline of the same aliquot using Spearman correlation. Similar to what we have described before for mRNA expression, only 9516 shared aliquots, and 641 shared mature miRNAs are used in this comparison. As shown in the boxplots at the bottom of Figure 5, the majority of the samples have a correlation coefficient of greater than 0.975, with an average of 0.984. Three cancer types, COAD (Colon adenocarcinoma), READ (Rectum adenocarcinoma) and OV (Ovarian carcinoma), have relatively lower correlation, which may reflect specific sensitivities of miRNA species in these cancer types to the reference genome and miRNA database versions.

**Figure 5.**
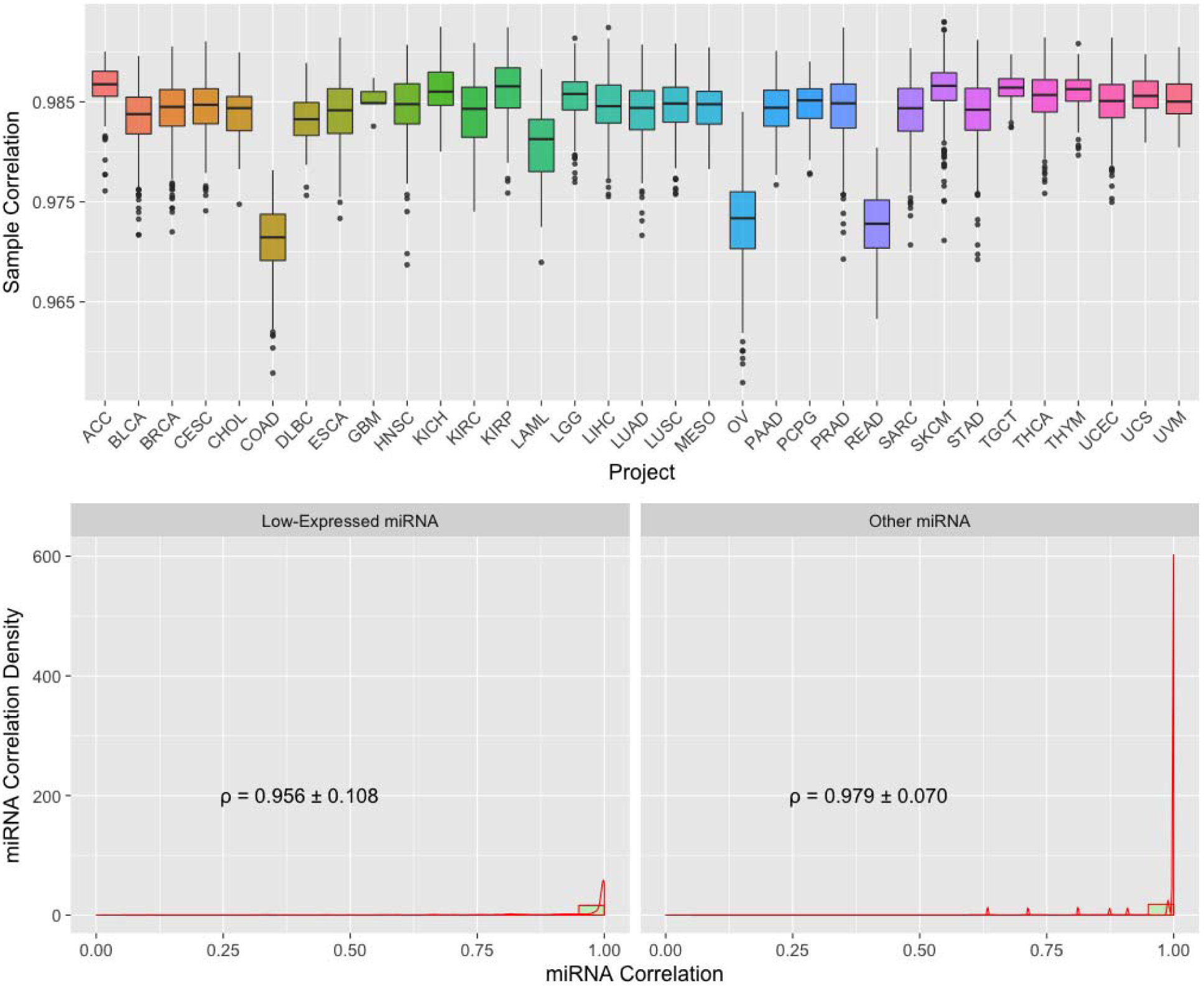
Boxplots of Spearman Correlation between GDC and TCGA miRNA Expression. Top: Boxplots of Sample to Sample Correlation between GDC and TCGA by Project. Bottom: Combined Boxplots and Density Plots of miRNA to miRNA Correlation between GDC and TCGA by Average Expression Level. All miRNAs are categorized in “Low-Expressed” and “Other” groups. Mean and standard deviation of miRNA to miRNA Spearman’s correlations are shown.

Expression level groups are also created in the miRNA datasets for detailed correlation comparisons (Figure 5. Bottom). Because there are many fewer miRNAs compared to mRNA genes, and many of them are expressed at low expression levels, we categorized miRNA into two groups. In the “Low-Expressed” group, all miRNAs show low (Q1 or Q2) in both TCGA and GDC quantifications; and the rest of miRNAs belong to the “Other” group. The average Spearman correlation in these two groups are about 95.6% and 97.9%, respectively.

### 7. Array-Based Copy Number Variation Data Processing

TCGA used Affymetrix Genome-Wide Human SNP 6.0 (SNP6) array data to identify genomic regions that have Copy Number Variations (CNV) by aggregation of typed loci into larger contiguous regions. Direct liftover of region boundaries from hg19 to GRCh38 results in fragmented segments with poor data quality due to change of probe loci between different genome builds. For this reason, the GDC SNP6 pipeline is built onto the existing TCGA level 2 tangent-normalized copy number data generated by Birdsuite ^31^ and uses the DNAcopy version 1.44.0 ^32^ R-package to perform a Circular Binary Segmentation (CBS) analysis ^33^ with GRCh38 probeset metadata. This pipeline normalizes noisy intensity measurements into chromosomal regions of equal copy number in the form of segment mean values, which are equal to log2(copy-number/2), so that diploid regions will have a segment mean of zero, amplified regions will have positive values, and deletions will have negative values.

To be consistent with the original TCGA data, two different output files were produced for each input: copy number segment files that generated from all probes, and masked copy number segment files, equivalent to the original TCGA “nocnv” file, generated by excluding certain probes that have been previously identified to carry copy number variations in a pool of germline samples.

### 8. Array-based Methylation Data Processing

TCGA used Illumina Infinium HumanMethylation27 (HM27) and HumanMethylation450 (HM450) BeadChip to measure the level of methylation at known CpG sites as beta values, calculated from array intensities (Level 2 data) as Beta = M/(M+U), where M is the methylated probe intensity and U is the unmethylated probe intensity. The GDC inherited these beta values from existing hg19-based TCGA Level 3 DNA methylation data, and re-annotated each probeset with new metadata information based on GRCh38 and GENCODE v22.

### 9. Integrated Genomic Data Clustering

To show how users can take advantages of GDC harmonized datasets for cross-project analysis, here we demonstrate integrated analysis with t-Distributed Stochastic Neighbor Embedding (t-SNE) ^34,35^ to reduce high dimensional information in the molecular data to two dimensions. (Figure 6). We utilized four distinct GDC generated molecular features of TCGA primary tumor samples, including mRNA expression, miRNA expression, methylation and copy number variation. Our results are shown only for TCGA data, but could be expanded to other GDC projects.

**Figure 6.**
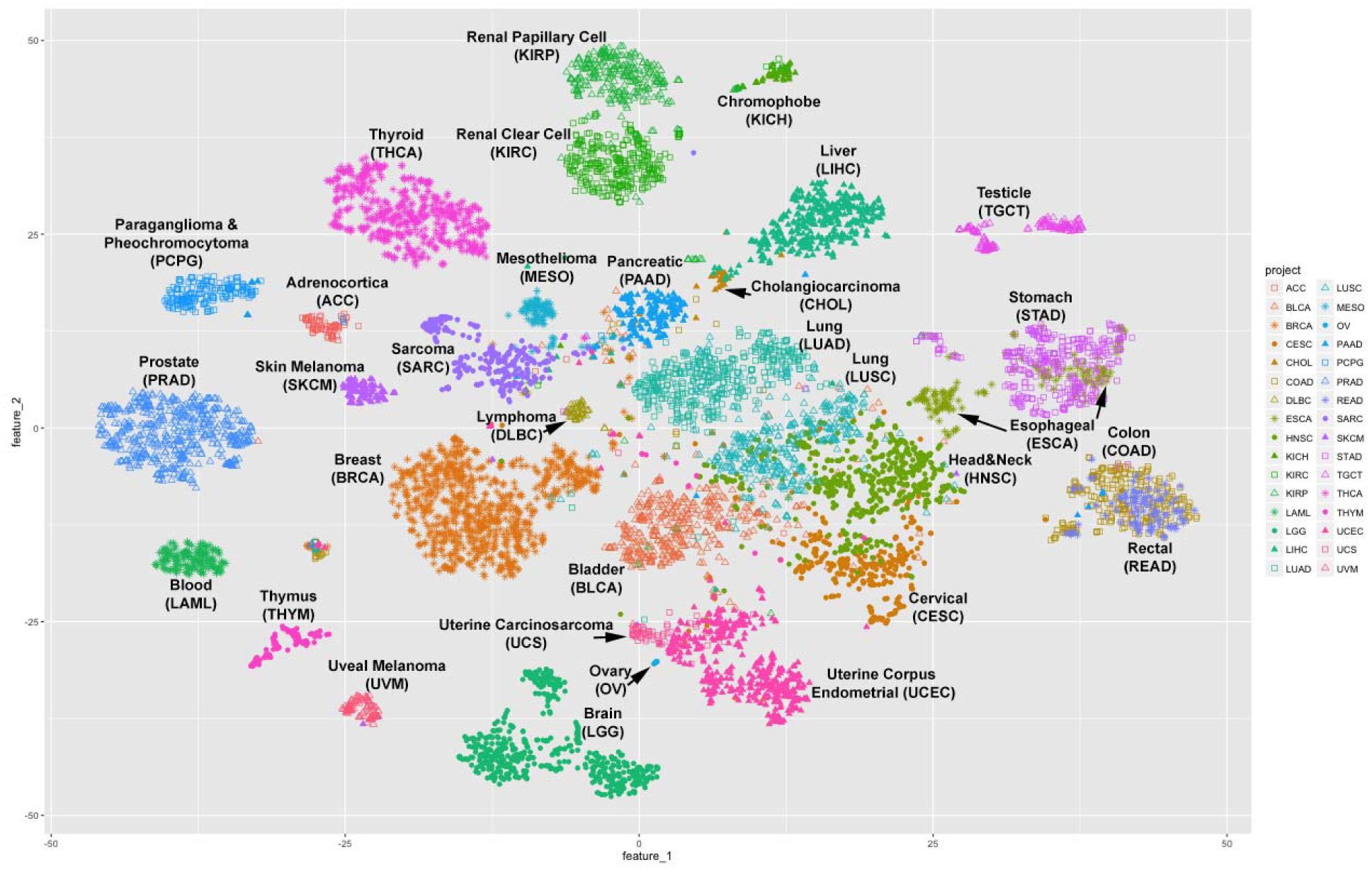
2D t-SNE Clustering of 32 TCGA Projects. Patients in different projects are presented by different dot colors or shapes.

In our result, different cancer types are well separated and some interesting patterns arise. For example, it has been previously suggested that colon and rectum cancer should be grouped as one colorectal cancer ^36^ and that esophageal adenocarcinomas resembles a subtype of stomach cancer ^37^. Our method supports those observations. In our 2D-clustering the majority of the colon cancer (COAD) samples are co-clustered with rectum cancer (READ) samples, and some esophageal cancer (ESCA) samples cluster together with stomach cancer (STAD) samples. We also identify a few samples designated as primary tumor in one cancer type, but which cluster together with samples from another cancer type. For example, a paraganglioma and phenochromocytoma (PCPG) sample is located within a cluster of adrenocortical carcinoma (ACC) samples.

## Conclusion

The rapid decrease in sequencing costs has lead to a rapid increase in the resources needed for storage and computation. The GDC provides a solution for this problem by centralizing storage and processing of genomics data. This model currently enables researchers to perform analysis in three different ways: 1) Quick data analysis and exploration using existing GDC visualization tools without the need to download files; 2) Data analysis using GDC generated high level processed data. These files are much smaller than the raw data; 3) Resource-extensive data analysis by downloading raw sequencing data to other data centers or commercial clouds.

The genomic data available through the GDC are analyzed uniformly using common algorithms and pipelines, and will be reanalyzed in the future as improved algorithms and methods are developed. As there is no consensus among the scientific community on the best algorithms on somatic variant detection, the GDC implements four callers, and each generates its own set of variant calling output. As shown in Figure 2A and 2B, only 55.97% of the variants are called by all four pipelines; while this category also includes 71.57% of the TCGA validated variants. This suggests a simple strategy to combine results from multiple callers to increase specificity. The boxplots in the bottom of Figure 3 show the effect of a combined caller approach to the recovery rate of TCGA validated variants by different cancer types. The downside of using a consensus call is a decrease in sensitivity. While we cannot measure sensitivity directly, a good approach may be to combine results from only two somatic callers rather than require a variant be called by all four pipelines. The GDC is in the development stage to generate a new set of MAFs that contain merged results from each tool.

The GDC data harmonization process also makes cross-project analysis much simpler, and reduces artificial findings due to differences in algorithms. Users will be then able to perform joint analysis of data originating from different projects because all data are processed by the same tools and represented in the same format.

The NCI’s Genomic Data Commons has provided a feasible and scalable model for easy sharing of large set of genomics data. We hope our efforts in data sharing and uniform data processing will be valuable not only to researchers, but also to clinicians, patients, and other interested parties, and help accelerate the long-term goal of precision oncology.

## Materials and Methods

### 1. Pipeline Development and Production

The GDC takes advantage of containerization technology and built pipelines using Docker (www.docker.com) to ensure pipeline scalability, portability, and reproducibility. The alignment was managed by an in-house job management system that creates new virtual machines of the designated configuration on demand in an OpenStack environment. The corresponding Docker container that holds the entire workflow then runs inside of the virtual machine for an additional layer of security.

During development of high-level data generation pipelines, we transitioned from single Docker workflows to using Common Workflow Language (CWL, www.commonwl.org) to describe analysis workflows of multiple Dockerized tools. CWL provides an additional transparent layer between workflow description and workflow execution, that allows even better scalability through parallel execution and portability. The GDC managed CWL pipeline production uses the Slurm Workload Manager (slurm.schedmd.com).

### 2. DNA-Seq Alignment and Co-cleaning

The GDC DNA-Seq alignment pipeline follows GATK Best Practices (https://software.broadinstitute.org/gatk/best-practices/). The main steps include regenerating FASTQ files from BAM input on a per-read group basis using biobambam2 ^38^ and alignment by read group using BWA (version 0.7.12) in both paired-end and single-end mode. This was followed by BAM sort, merge, and MarkDuplicates using Picard ^39,40^. The GDC maps reads using either BWA MEM mode if length is equal or larger than 70bp or BWA Aln mode if below. BWA Aln is also used when mapping older FASTQ reads formatted with Illumina-1.3 and Illumina-1.5 quality scores. Multiple QC metrics were collected both before and after realignment using FastQC ^41^, samtools ^39^ and Picard. Re-aligned BAM files from the same patient are then collected together for co-cleaning using GATK 3.6 IndelRealigner and BaseQualityScoreRecalibration.

The TCGA and TARGET BAMs to be harmonized were originally processed over a relatively large time scale in relation to the development of NGS technology. Some originated from as early as 2010 and were generated by a variety of workflows and reference genomes (11 different reference genomes in total including variants of hg18, hg19, GRCh36 and GRCh37). Some of the workflows had introduced incorrect information into the data which were corrected in the GDC harmonization process.

### 3. MuTect2 Somatic Variant Calling

MuTect2 is built upon the capability of local de novo assembly by HaplotypeCaller and somatic genotyping engine of Mutect. Mutect applies a Bayesian classifier to detect somatic mutations^11^. The GDC uses MuTect2 tools from the GATK nightly-2016-02-25-gf39d340 version. Before tumor normal pairs can be used for somatic variant calling, it is important to generate a Panel of Normals (PoNs) filter that contains calling artifacts and potential germline variants. As mentioned previously, whole genome amplified (WGA) samples are analyzed with dontUseSoftClippedBases turned on.

### 4. VarScan2 Somatic Variant Calling

VarScan2 is another somatic variant caller that identifies both SNV and INDELS. It uses heuristics and statistics to identify variants and considers the confounding impacts of read depth, base quality, variant allele frequency and statistical significance ^13^. GDC uses VarScan2 version 2.3.9.

The first step of VarScan2 calling is to generate a mpileup file of both tumor and normal BAMs using samtools for a single mpileup file. We set the quality cutoff for samtools to be 1 and also disabled Base Alignment Quality (BAQ) score computation. The mpileup is then used as input to VarScan Somatic to generate a VCF file that contains both SNP and INDEL calls. The resulting VCF is filtered for significant calls using VarScan ProcessSomatic.

### 5. MuSE Somatic Variant Calling

MuSE calls somatic variants using Markov Substitution model for Evolution ^12^. The first step, “MuSE call”, estimates the equilibrium frequencies of all four alleles and presents the maximum *a posteriori* on every genomics locus. The second step, “MuSE sump”, performs a tier based cutoff based on a sample-specific error model which also takes dbSNP information into account. GDC uses MuSE version 1.0rc_submission_c039ffa.

Parallelization can be implemented for the first step of MuSE, based on genomic chunks, which can accelerate the production close to linear. The GDC currently only passes calls with quality filter “PASS” to the GDC public MAF files; however, variants with other quality Tier values could also be considered a user’s discretion.

### 6. SomaticSniper Somatic Variant Calling

SomaticSniper is a somatic variant caller that only identifies SNPs. It uses a bayesian inference to compare genotype likelihoods between tumor and normals ^14^. GDC uses the default parameter settings of SomaticSniper version 1.0.5.0.

### 7. Somatic Variant Filters

In addition to the built-in filters in each somatic caller, the GDC also applies additional filtering tools to label caller-generated variants. Because these filters are frequently updated, we have highlighted only a few of the major steps below.

A. False Positive Filter (FPFilter, https://github.com/ucscCancer/fpfilter-tool) was applied to both VarScan2 and SomaticSniper VCFs.
B. SomaticSniper variants with SSC < 25 are removed from annotated VCFs. This is the only step in the entire GDC somatic variant pipeline in which low-quality variants are removed, instead of tagged.
C. A WXS Panel of Normals were generated internally by MuTect2 calling on about 5,000 TCGA normal WXS samples in artifact detection mode, and combined using GATK CombineVariants. This is not only used as a MuTect2 built-in filter ^42^, but also applied to the other three somatic calling outputs in a similar manner.
D. d-ToxoG (http://archive.broadinstitute.org/cancer/cga/dtoxog) is used to remove oxoG artifacts from point mutation calls. These artifacts were generated due to oxidative DNA damage during sample preparation ^43^.
E. DKFZ Strandbias Filter (https://github.com/eilslabs/DKFZBiasFilter) is used to tag variants that are supported with significant bias from one strand direction compared to the other.

### 8. MAF Generation

Mutation Annotation Format (MAF) is a tab-delimited text file with aggregated mutation information from VCF Files and are generated on a project-level. The GDC currently produces two types of MAF files: controlled-access MAFs that contain all variants in VCFs, and open-access somatic MAFs that contain “high quality” variants and reduced germline contaminations. Any user can explore the open-access somatic MAF for high quality calls; while a more sophisticated user may want to apply for dbGaP access to obtain the superset of mutations in the controlled-access MAF. With the larger set of mutations they may perform custom filtering based on FILTER and GDC_FILTER columns, or collect information that was removed from the open-access version, such as supporting read depth in the normal samples.

The specification of the GDC MAF can be found at https://docs.gdc.cancer.gov/Data/File_Formats/MAF_Format/.

### 9. RNA-Seq Alignment

The RNA-Seq alignment pipeline performs alignments of raw reads against the reference genome using a two-pass approach using STAR ^23,43^. The first pass alignment recognizes splice junctions in the sample, and the second pass uses those splice junctions to perform the final alignment. STAR version 2.4.0f1 was initially used and we then switched to version 2.4.2a that fixed a bug and allowed us to complete processing all the input files. If both BAM and FASTQ input files existed, only FASTQ files were used.

### 10. HTSeq mRNA Quantification

The HTSeq (version 0.6.p1) ^24^ pipeline is used for calculating the number of reads that align to different genes in the genome. As mentioned previously, only reads that can be uniquely assigned to a gene are counted. The GDC ran HTSeq on all samples as unstranded libraries in order to maintain consistency for cross-sample comparisons.

The raw counts are normalized into Fragment per kilobase million mapped reads (FPKM) and Upper Quartile Normalized FPKM (FPKM-UQ) using all protein-coding genes as the denominator.

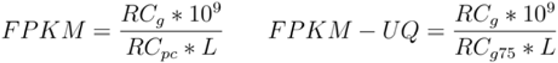

Where,

- **RCg:** Number of reads mapped to the gene
- **RCpc:** Number of reads mapped to all protein-coding genes
- **RCg75:** The 75th percentile read count value for genes in the sample
- **L:** Length of the gene in base pairs

### 11. DexSeq Exon Quantification

The GDC has also generated exon-level quantification using the DEXSeq ^44^ pipeline. The first step in this pipeline is to create the flattened General Feature Format (GFF) file, which essentially collapses the information for multiple transcripts spanning the same exon into exon counting bins for that exon. Once the flattened GFF file is obtained, the number of reads that overlap with each exon counting bin are calculated. The result is a flat file which has raw counts for each exon. This data type is not currently available in the GDC data portal and will appear in a later data release.

### 12. miRNA-Seq Alignment

The GDC miRNA harmonization pipeline begins with a realignment of TCGA and TARGET miRNA-Seq reads using a similar strategy of the GDC DNA-Seq alignment pipeline. Because reads of miRNA-Seq are typically short, only BWA Aln was used.

### 13. miRNA Profiling

miRNA quantification is done with a modified version of the miRNA Profiling Pipeline v0.2.7 ^30^ from BCGSC (British Columbia Genome Sequencing Center). In this pipeline, miRNA species and miRNA isoforms are counted differently, and normalized Reads Per Million (RPM) values are also derived. The final results from each miRNA-Seq sample is a miRNA species quantification file and a miRNA isoform quantification file, in a human-readable format compatible to the original TCGA data.

### 14. SNP 6.0 Array Copy Number Segmentation

The hg19-based probeset metadata were obtained from the Affymetrix website, and then lifted over to GRCh38. Probes with reference bases not matching between hg19 and GRCh38 were removed.

To generate Copy Number Segment file, all SNP and CNV probes are used for Circular Binary Segmentation (CBS) calculation, with the only exception that probes in the Pseudo-Autosomal (PAR) regions were removed in males prior to calculation. To generate the Masked Copy Number Segment file from this result, all probesets in chromosome Y and in the frequent copy number variant regions in germlines obtained from GenePattern were also removed prior to calculation.

### 15. Methylation Array Beta Value Annotation

Using probe sequence information provided in the manufacturer’s manifest, HM27 and HM450 probes were remapped to the GRCh38 reference genome ^45^. Type II probes with a mapping quality of <10, or Type I probes for which the methylated and unmethylated probes map to different locations in the genome, and/or had a mapping quality of <10, had an entry of ‘*’ for the ‘chr’ field, and ‘-1’ for coordinates ^45^. These coordinates were then used to identify the associated transcripts from GENCODE v22, the associated CpG island (CGI), and the CpG sites’ distance from each of these features. Multiple transcripts overlapping the target CpG were separated with semicolons. Beta values were inherited from existing TCGA Level 3 DNA methylation data (hg19-based) based on Probe IDs.

### 16. Variant Comparison

The same genetic variant can be represented in VCF format in multiple different ways ^46^, and many of these discrepancies can not be easily solved by existing normalization tools. In order to reduce false-positive annotations, GDC requires a strict matching of Chromosome, Position and Alternative Alleles during implementation of MAF annotations. However, in various variant comparisons in this paper, we applied a loose matching strategy to regard two variants the same if they have overlapping regions between starting and ending positions. This is particularly useful when a non-INDEL caller, such as SomaticSniper or MuSE, represents a INDEL site as point mutations.

### 17. t-SNE Clustering

mRNA expression count, miRNA expression count, Copy Number Segmentation, and Methylation Beta Values were collected from the GDC Data Portal. For mRNA expression and miRNA expression, we removed low-expressed genes and miRNAs if 99% or more samples have less than or equal to 1 count. For Copy Number Segments, we computed average segmentation means on each gene weighted on overlapped length between segments and genes. For methylation data, we removed probes that are empty in more than 5% of the samples, and imputed the remaining empty values with the probeset mean. The methylation data is also randomly down-sampled by 25% of the probes to reduce computational burden.

To integrate these data together for t-distributed stochastic neighbor embedding (t-SNE) clustering, we assigned the following arbitrary weights on these four data types (3:3:1:1 for mRNA:Methylation:miRNA:CNV). Then we used Principal Component Analysis (PCA) to generate the top 200 Principal Components (PCs) with the R function prcomp, and ranked these 800 PCs by weighted “variation explained”. The top 200 PCs out of this analysis were scaled again to match the desired weights among these data types, which were used as input for t-SNE analysis with the R package Rtsne. We ran t-SNE 1000 times with random seeds, and displayed the result that minimized “cost”.

## Acknowledgements

The authors would like to acknowledge the following individuals for consultations in pipeline development: Kyle Ellrott, Cyriac Kandoth, Gordon Saksena, Gad Getz and Li Ding of the TCGA multi-center mutation calling in multiple cancers Calling 3 (MC3) Working Group (reference to the paper when it publishes); Angela Brooks, Nuno Fonseca and Andre Kahles of the PanCancer Analysis of Whole Genomes (PCAWG) RNA-Seq Working Group; Yussanne Ma, Denise Brooks and Gordon Robertson of Canada’s Michael Smith Genome Sciences Centre; Wanding Zhou and Hui Shen of the Van Andel Institute; and Yu Fan and Wenyi Wang of the University of Texas MD Anderson Cancer.

This project has been funded in in part with Federal funds from the National Cancer Institute, National Institutes of Health, Task Order No. 17X053 and Task Order No. 14X050 under Contract No. HHSN261200800001E. The content of this publication does not necessarily reflect the views or policies of the Department of Health and Human Services, nor does mention of trade names, commercial products, or organizations imply endorsement by the U.S. Government.

